# Inhibition of *Neisseria gonorrhoeae* complement-mediated killing during acute gonorrhoea is dependent upon the IgG2:IgG3 antibody ratio

**DOI:** 10.1101/2023.09.26.558794

**Authors:** Samantha A. McKeand, Sian E. Faustini, Alex Cook, Nikki Kennett, Mark T. Drayson, Adam F. Cunningham, Ian R. Henderson, Jonathan D. C. Ross, Jeffrey A. Cole, Amanda E. Rossiter-Pearson

## Abstract

Excessive binding of antibodies to the bacterial cell surface can paradoxically increase resistance of some Gram-negative pathogens to complement-mediated killing (CMK). We examined CMK of 336 *Neisseria gonorrhoeae* clinical isolates sampled from participants recruited to a clinical trial. Serum bactericidal assays revealed 3% (9/336) of the autologous participant sera that were tested inhibited CMK. Gonococci isolated from these participants were resistant to the autologous host serum, sensitive to a pool of healthy control sera (HCS) and protected by the host serum in a 1:1 mixture with HCS. Analysis of the clinical metadata showed that there were a significantly higher proportion of inhibitory sera found in participants with urethral infections and from men within the transmission network of men who have sex with women (MSW), when compared to the whole cohort. Following antibody purification from selected participants with inhibitory sera (5/9), IgG and IgM protected the autologous isolates from HCS-mediated killing. Only three of these isolates were protected by purified IgA. A closer examination of IgG subclasses using whole gonococcal cell ELISAs revealed a strong correlation between increased IgG2 binding and decreased IgG3 binding to the bacterial cell surface of isolates that were resistant to CMK. This suggests that IgG2 prevents bactericidal IgG3 from initiating CMK and that the IgG2:IgG3 ratio is important for determining either inhibition or killing of isolates. We therefore reveal a previously unreported mechanism by which inhibitory antibodies prevent CMK of *N. gonorrhoeae*.

## Introduction

Gonorrhoea is a sexually transmitted infection (STI) caused by the Gram-negative bacterium *Neisseria gonorrhoeae*. It is the second most common bacterial STI in England, with 82,592 diagnosed infections reported in England in 2022 (UKHSA, 2023) representing a 50.3% increase in infections from the previous year. Gonococcal disease is typically an uncomplicated infection, which infects the genitourinary tract of both women and men, as well as mucosal surfaces of the oropharynx and the rectum. Although symptoms are more common in men than in women, disease complications include epididymo-orchitis in men and pelvic inflammatory disease and infertility in women (Lenz and Dillard, 2018, Unemo *et al*., 2019). Disseminated gonococcal infection (DGI) is a severe complication involving a systemic form of the disease that can occur in 0.5-3% of individuals (Beatrous *et al*., 2017).

The emergence of antimicrobial resistance in *N. gonorrhoeae* raises the concern that untreatable gonorrhoea might be inevitable (Unemo and Nicholas, 2012). Therefore, there is an urgent requirement to develop an effective vaccine. Despite many efforts spanning a century, there is currently no FDA-approved vaccine available for gonorrhoea (Russell *et al*., 2019). However, surveillance data following a mass vaccination campaign in New Zealand with an outer membrane vesicle (OMV)-based vaccine against meningococcal B (MeNZB) revealed an association with reduced rates of gonorrhoea in young adults (Petousis-Harris *et al*., 2017). These observations indicated that protection can be induced against *N. gonorrhoeae*, and have motivated ambitions to produce an OMV-based vaccine specific for *N. gonorrhoeae*.

Antibodies typically protect the host against pathogens. However, there have been some instances in which antibodies have been directly linked with the exacerbation of bacterial disease, as detailed in a review by Torres *et al*. (2021). These inhibitory antibodies were identified in the serum of patients with Gram-negative infections, including *Salmonella enterica* serovar Typhimurium, *Pseudomonas aeruginosa* and *Escherichia coli* (MacLennan *et al*., 2010, Wells *et al*., 2014, Goh *et al*., 2016, Coggon *et al*., 2018, Pham *et al*., 2021). Such patients were found to have IgG2, IgM or IgA antibodies specific to the O-antigen component of LPS, with significantly higher antibody titers or greater antibody affinity than those observed in non-infected individuals. The proposed mechanisms for this phenomenon involve these antibodies leading to the deposition of complement away from the bacterial membrane, or creating an antibody “blockade” so that access for bactericidal antibodies is obstructed. Moreover, symptoms of patients with *P. aeruginosa* infections and severe bronchiectasis improved when inhibitory antibodies were removed by plasmapheresis (Wells *et al*., 2017). In addition, earlier studies characterised inhibitory antibodies against *N. gonorrhoeae* that were isolated from normal healthy serum (McCutchan *et al*., 1978). These antibodies inhibited complement-mediated killing (CMK) of *N. gonorrhoeae* strains isolated from patients with DGI (McCutchan *et al*., 1978, Rice and Kasper, 1982). Removing these inhibitory antibodies *in vitro* restored CMK (Joiner *et al*., 1985, Rice *et al*., 1986). Further characterisation revealed that these antibodies were directed against the gonococcal outer membrane protein III, which is known as reduction modifiable protein (RmpM) (Rice *et al*., 1986). However, the nature and indeed the prevalence of these inhibitory antibodies were not characterised in these early studies. Most patients presenting with gonococcal infection in clinic are acute uncomplicated mucosal infections. There is limited research to understand whether these patients produce inhibitory antibodies during acute infection and, if so, their prevalence within this population and the implications for vaccine design against *N. gonorrhoeae*.

In this study, we utilised *N. gonorrhoeae* clinical isolates (336) and autologous serum from 283 participants collected as part of the G-ToG clinical trial (Ross *et al*., 2019). Using this set of samples, we aimed to determine whether inhibitory antibodies were produced by these infected participants and whether these were associated with any of the clinical metadata collected as part of the trial, such as gender, HIV status, sexual networks, anatomical site of infections and recurrent infections. We report that 3% of participants harboured serum that was able to protect their autologous isolates from CMK. Purified IgG, IgM and IgA antibodies from these sera were able to protect isolates from CMK by healthy control sera (HCS). However, protection was variable between the gonococci strains tested and we show that this variability was due to an increased IgG2 and decreased IgG3 binding to the bacterial cell surface. Ultimately, these data suggest that the mechanism underlying protection of *N. gonorrhoeae* from CMK is distinct to that of inhibitory antibodies against other Gram-negative bacteria, in which significantly higher titres of IgG2 alone protected bacteria from CMK. We propose that it is the IgG2:IgG3 ratio binding to the bacterial cell surface that determines survival of some gonococci in serum.

## Materials and methods

### Bacterial isolate collection and serum

From the 720 participants recruited to the G-ToG clinical trial (Ross *et al*., 2019), 336 *N. gonorrhoeae* isolates were collected from 283 individuals. *N. gonorrhoeae* were isolated from participants during 2014 (n=16), 2015 (n=126) and 2016 (n=141). Blood samples were also collected from participants and stored as serum by the Clinical Immunology Services (CIS) (The University of Birmingham, UK). Throughout this work, clinical isolates are named ‘NGxxx’, and serum isolated from the participant with the indicated isolate are referred to as ‘NGxxx-serum’. Upon trial entry, clinical and demographic data were also collected from participants and recorded in a database developed by the Nottingham Clinical Trial Unit (NCTU) within the G-ToG team. Separately, blood was taken from eight healthy human volunteers with no previous history of gonococcal infection, where equal amounts of each HCS sample were pooled and stored at -80°C.

### Growth conditions

Bacteria were routinely grown on GC agar plates supplemented with Vitox (Oxoid) in a candle extinction jar for 20 hours at 37°C, and then single colonies were passaged onto fresh plates and grown for another 16-18 hours.

### Serum bactericidal assay

A serum bactericidal assay (SBA) was used to determine the susceptibility of gonococcal isolates to CMK. Bacteria from day two plates were resuspended in PBS and cultures were incubated at 37°C with the indicated serum at a final concentration of 1.5 x 10^7^ colony forming units (CFUs) per ml. As a positive control, the same cultures were added to PBS and incubated under the same conditions. Aliquots were taken before incubation and at hourly intervals for 3 hours and plated in triplicate onto GC agar. After overnight growth at 37°C, survival was calculated from the number of viable colonies at selected time points relative to the baseline colony counts at 0 h.

For SBAs involving purified antibody, bacteria were prepared in the same way, and added to a mixture of 50% (v/v) HCS and individual purified antibody at the indicated concentrations in PBS. When an exogenous complement source was required, bacteria were added to a mixture of 5% baby rabbit complement (BRC) (AbD Serotec) and individual purified antibody at the indicated concentrations in PBS.

### Flow cytometric binding assay

Binding of antibodies IgG, IgM and IgA to isolates was measured using flow cytometry. Bacteria from day two plates were resuspended in PBS and cultures were incubated with autologous host serum at a final concentration of 2 x 10^8^ CFU/ml and incubated without aeration at 37°C for 20 min. Fluorochromes DyLight 405, PerCP-Cy5.5 and APC-Cy7 were conjugated to anti-human murine monoclonal total IgG (clone #IgG-R10) (Lund *et al*., 1996), IgM (clone IgM-AF6) (Grafton *et al*., 1997) and IgA (clone # IgA-2D7) (Mestecky *et al*., 1996), respectively using Lightning-Link® Rapid Conjugation kits (Innova Biosciences, Cambridge, UK). Each secondary antibody was added to each sample at a concentration of 0.5 mg/ml, bacteria were fixed and then re-suspended in cellWASH (BD Biosciences) for flow cytometric analysis. Data were collected on the FACSCanto II (BD Biosciences, Oxford, UK) using FACSDiva (BD Biosciences) software, and data were then analysed using FlowJo software (BD Biosciences).

### Antibody purification

Antibody isotypes IgG, IgM and IgA were each individually depleted and purified from 500 µl of participant sera. In brief, individual serum samples were first run through a Protein G column at 1 ml/min and purified IgG was then eluted at 2 ml/min in a fraction size of 5 ml. The unbound serum (depleted of IgG) was then used as the start material for depletion of IgA through a Capture Select IgA column. The unbound serum (depleted of IgG and IgA) was then run through a Capture Select IgM column to purify IgM.

### Enzyme-linked immunosorbent assay

An enzyme-linked immunosorbent assay (ELISA) was used to determine IgG subclasses binding to gonococcal isolates. Bacteria from day two plates were resuspended in PBS and adjusted to an OD 600 _nm_ of 0.5 to coat high binding ELISA plates (Nunc-Immuno™, ThermoScientific). After incubation overnight at 4ºC, plates were blocked with 1% bovine serum albumin (BSA) in PBS for 1 hour. Participant sera pre-diluted to 1 in 20 were added to plates and serially diluted 3-fold in dilution buffer (0.05% Tween20 and 1% BSA in PBS). After incubation for 1 hour at 37ºC, plates were washed with wash buffer (0.05% Tween 20 in PBS). Next, horseradish peroxide (HRP) conjugated secondary antibodies for measurement of total IgG (SouthernBiotech) and individual IgG subclasses 1-4 (SouthernBiotech) were diluted to manufacturer recommended concentrations and incubated for 1 hour. The plates were developed using TMB-core (AbD Serotec), and an OD 450 _nm_ of each well was read using a CLARIOstar (BMG Labtech).

### Statistical analysis

Prism version 9.2.0 (GraphPad software, CA) was used for a Mann-Whitney U statistical test of significance for analysis of flow cytometry data and an unpaired t-test of significance when analysing ELISA data. For the comparison of clinical data between the participants with inhibitory sera and the total population, a two proportion z-test with a 95% confidence level was applied, with the resultant z-score used to calculate statistical significance using a 2-tailed z-test. All statistical analysis throughout was represented as * (*p*≤0.05), ** (*p*≤0.01), *** (*p*≤0.001) and **** (*p*≤0.0001).

## Results

### Screening a collection of clinical Neisseria gonorrhoeae isolates for resistance to autologous host serum

To determine whether *N. gonorrhoeae* induced inhibitory antibody production, we examined the collection of clinical isolates (n=336) from participants (n=283) with acute gonorrhoea infection obtained as part of the G-ToG clinical trial (Ross *et al*., 2019). Each isolate was screened for resistance to CMK by incubating bacteria in serum of the host from which it was isolated. As expected, 84% (282/336) of the gonococcal isolates were killed by the host serum, demonstrated by a log_10_ kill of greater than 8 after 3 hours. A representative example of an isolate that was sensitive and resistant to CMK is shown in Fig. 1A and B, respectively. One reason for resistance to serum could be that the complement is inactive due to blood processing. To rule this out and standardise the study of antibody, an exogenous complement source, BRC, was added to heat-inactivated participant sera in similar experiments to eliminate these sera. This assay identified eight isolates that were killed by BRC and therefore only appeared resistant because the human complement component had not been able to mediate cell lysis. These eight isolates were not studied further. Therefore, we identified 46 isolates that were resistant to CMK, indicating that the participant autologous serum might contain inhibitory antibodies (Fig. 1C).

**Figure 1.**
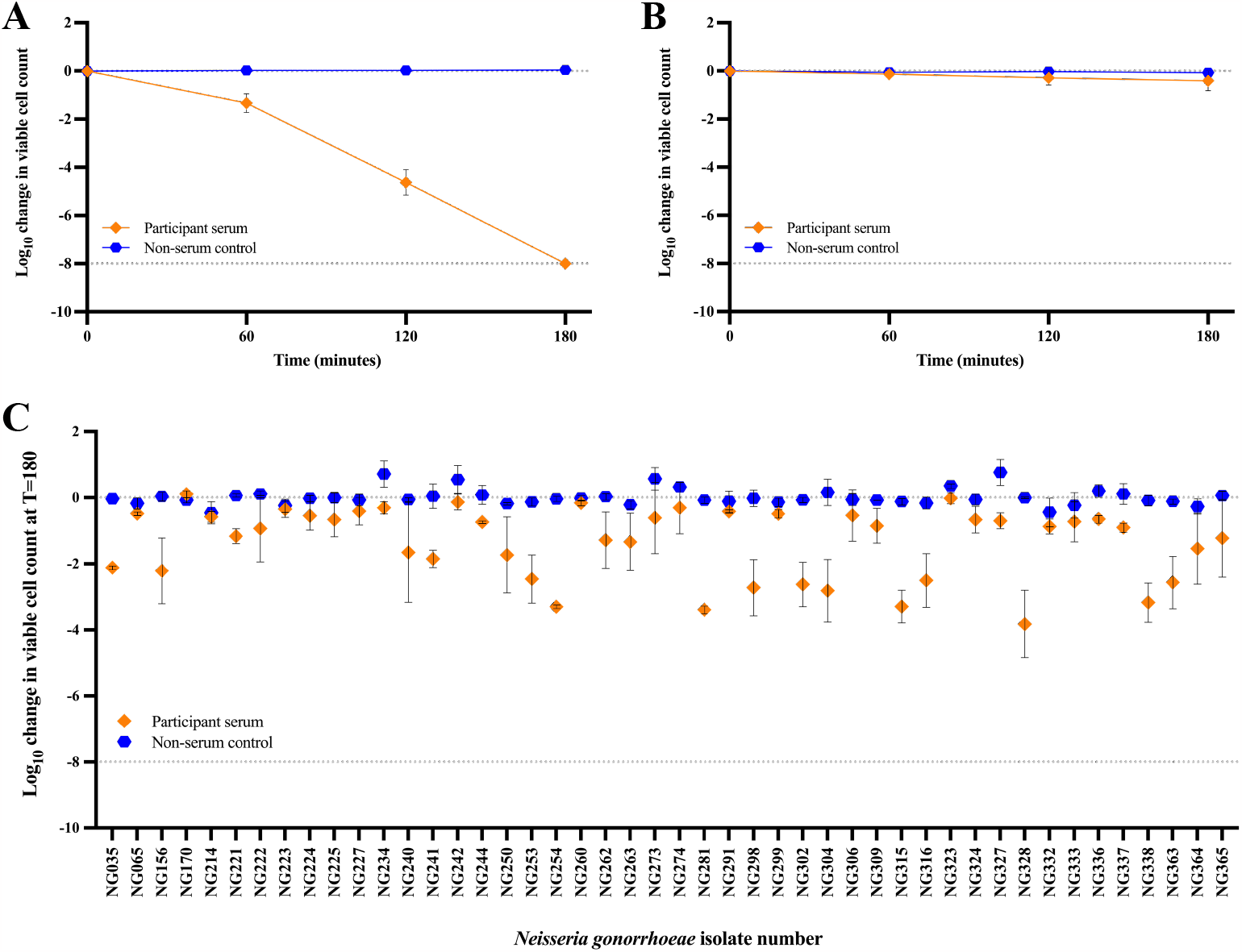
Screening the *Neisseria gonorrhoeae* clinical isolates for resistance to their autologous host serum. A serum bactericidal assay (SBA) was used to screen the 336 *N. gonorrhoeae* clinical isolates to determine their sensitivity to autologous participant serum. Cultures of each isolate were incubated with either the autologous serum, or PBS as a non-serum control. The change in viable colony forming units for **A)** isolate NG161, a representative isolate that was sensitive to killing by autologous serum and **B)** isolate NG227, a representative isolate that was resistant to killing by autologous serum, were determined after 60, 120 and 180 min, with negative values corresponding to a decrease in viable bacteria when compared to the initial inoculum. Values at -8 indicate the limit of detection for samples from which no colonies formed. **C)** The change in viable colony forming units after 180 min in autologous serum for the 46 *N. gonorrhoeae* clinical isolates that were still viable after SBA completion. Data shown are the mean and standard deviation from three independent experiments.

### Effects of pooled healthy serum on isolates that resisted killing by autologous host serum

Some of the 46 isolates might have survived incubation with the host serum because they are intrinsically resistant to CMK, independent of specific antibodies. Such isolates should also be resistant to killing by pooled HCS from uninfected individuals with no known previous history of gonococcal infection. Serum bactericidal assays were therefore repeated using pooled HCS. The 46 isolates screened fell into two groups: 26 of the isolates resisted killing by HCS, which suggests that these isolates are potentially intrinsically resistant to serum killing. The remaining 20 isolates were completely killed by HCS (Fig. 2A). As these 20 isolates are sensitive to HCS-mediated killing, but resistant to killing in autologous sera, the sera from these participants merited further investigation for the presence of inhibitory antibodies.

**Figure 2.**
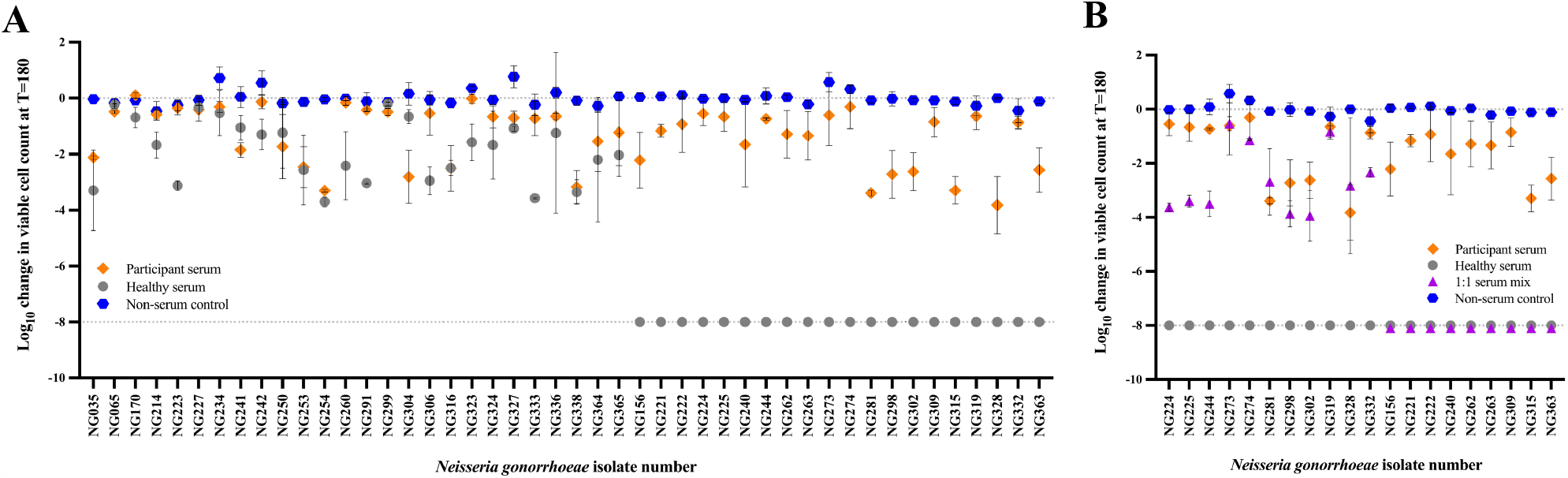
Effects of healthy sera on *Neisseria gonorrhoeae* isolates that resisted killing by their autologous host serum. A serum bactericidal assay was used to screen **A)** the 46 *N. gonorrhoeae* clinical isolates that resisted killing by their autologous participant serum, to determine the effect of healthy control serum (HCS) on survival. Cultures of each isolate were incubated with either the autologous serum, HCS or PBS as a non-serum control. **B)** The resultant 20 *N. gonorrhoeae* clinical isolates that succumbed to killing by HCS, were assayed to determine the effect of a 1:1 mix of both sera on survival. Cultures of each isolate were incubated with either the autologous serum, HCS, 1:1 mix of both sera or PBS as a non-serum control. For all experiments, the change in viable colony forming units after 180 min in each sera for each *N. gonorrhoeae* isolate was measured, with negative values corresponding to a decrease in viable bacteria when compared to the initial inoculum. Values at -8 indicate the limit of detection for samples from which no colonies formed. Data shown are the mean and standard deviation from three independent experiments.

Inhibitory antibodies are typically defined by their ability to prevent HCS-mediated killing. Thus, each of the 20 isolates were incubated with a 1:1 mixture of HCS and the autologous participant serum (Fig. 2B). Nine of the 20 isolates were killed by the mixed sera, suggesting that the participants had failed to generate an effective immune response against their autologous isolate. However, eleven isolates were either completely or significantly more resistant to the mixed sera than to HCS. The autologous sera therefore protected these eleven isolates from HCS-mediated killing. These eleven isolates were isolated from nine of the 283 participants recruited to the trial.

Previous studies have linked the presence of inhibitory antibodies with clinical outcome such as HIV, bronchiectasis and urosepsis (MacLennan *et al*., 2010, Wells *et al*., 2014, Coggon *et al*., 2018). Therefore, we next sought to determine whether patients with protective sera were associated with any of the clinical or demographic metadata collected during recruitment to the trial. There were no significant differences in the proportions of participants with protective sera with ethnicity, gender, previous infections, HIV status and symptomatic or asymptomatic infections compared to the proportions within the whole cohort. However, there was a significant increase in the proportion of participants with protective sera in the men who have sex with women (MSW) sexual network (44% vs 21%) and participants with urethral infections (89% vs 55%) compared to the whole cohort (Supp. Table 1). Although outside the scope of this study, these associations provide the rationale for follow up studies to determine the clinical implications of protective sera on chronic and recurrent gonorrhoea in this subset of patients. Here, we focused our attention on the nine participants whose sera are strongest candidates for containing inhibitory antibodies.

### Effect of antibody in inhibitory sera on complement-mediated killing of Neisseria gonorrhoeae

A flow cytometry assay was set up to measure binding of antibodies IgG, IgM and IgA to the 46 resistant isolates and a set of 46 isolates that were sensitive to autologous host serum. Sensitive isolates chosen mirrored the participant demographic of the resistant isolates. There were no significant differences in the amount of binding of IgG or IgM from their autologous serum to the resistant isolates compared to the sensitive isolates (Supp. Fig. 1A and B). There was a small but significantly lower binding of IgA to resistant than to sensitive isolates (Supp. Fig. 1C). Given that these results were not consistent with previous studies showing that increased titers of antibody bound to bacteria (MacLennan *et al*., 2010, Wells *et al*., 2014, Goh *et al*., 2016, Coggon *et al*., 2018, Pham *et al*., 2021), we next assessed whether antibodies were the inhibitory factor in the participant serum that protected the gonococcal isolates from CMK. To do this, total IgG, IgM, and IgA were purified from the sera of five selected participants from the nine identified above, including NG244-serum, NG298-serum, NG302-serum, NG273/274-serum and NG332-serum. Note that isolates NG273 and NG274 were derived from the same participant at different anatomical sites. As a control, antibody preparations of total IgG, IgM and IgA were also purified from the serum of a participant able to kill their autologous isolate (NG115). Antibodies from NG115-serum retained their ability to elicit CMK and therefore were still functional after purification (data not shown). We next sought to test individual antibody isotypes from each of the five inhibitory sera for CMK. Thus, each isotype at varying concentrations was individually incubated with 50% HCS. The highest antibody concentrations tested (IgG at 1000 µg/ml, IgM at 100 µg/ml and IgA at 100 µg/ml) were all below normal physiological levels in the blood (6-16 mg/ml, 0.4-2.5 mg/ml and 0.8-3 mg/ml, respectively). As all selected isolates were sensitive to HCS-mediated killing, this modified assay would reveal whether the different antibody isotypes could inhibit CMK, and if so, at what concentration.

Only the highest concentrations of IgG (100-1000 µg/ml) and IgM (100 µg/ml) from autologous sera protected isolates NG244, NG298 and NG302 from HCS-mediated killing (Fig. 3A and B respectively). In contrast, matching purified IgA were not protective even at the highest concentration tested. Similarly, for NG273, only relatively high concentrations of IgG (100 µg/ml) and IgM (1 µg/ml) protected from HCS-mediated killing, but this isolate was protected by autologous IgA at concentrations above 1 µg/ml (Fig. 3C). Interestingly, isolates NG274 and NG332 were protected from HCS-mediated killing by much lower concentrations (from 0.0001 µg/ml) of all isotypes, IgG, IgM and IgA (Fig. 3A-C). This revealed that there are differences in how IgG is binding to autologous isolates or how IgG is preventing CMK in these participants.

**Figure 3.**
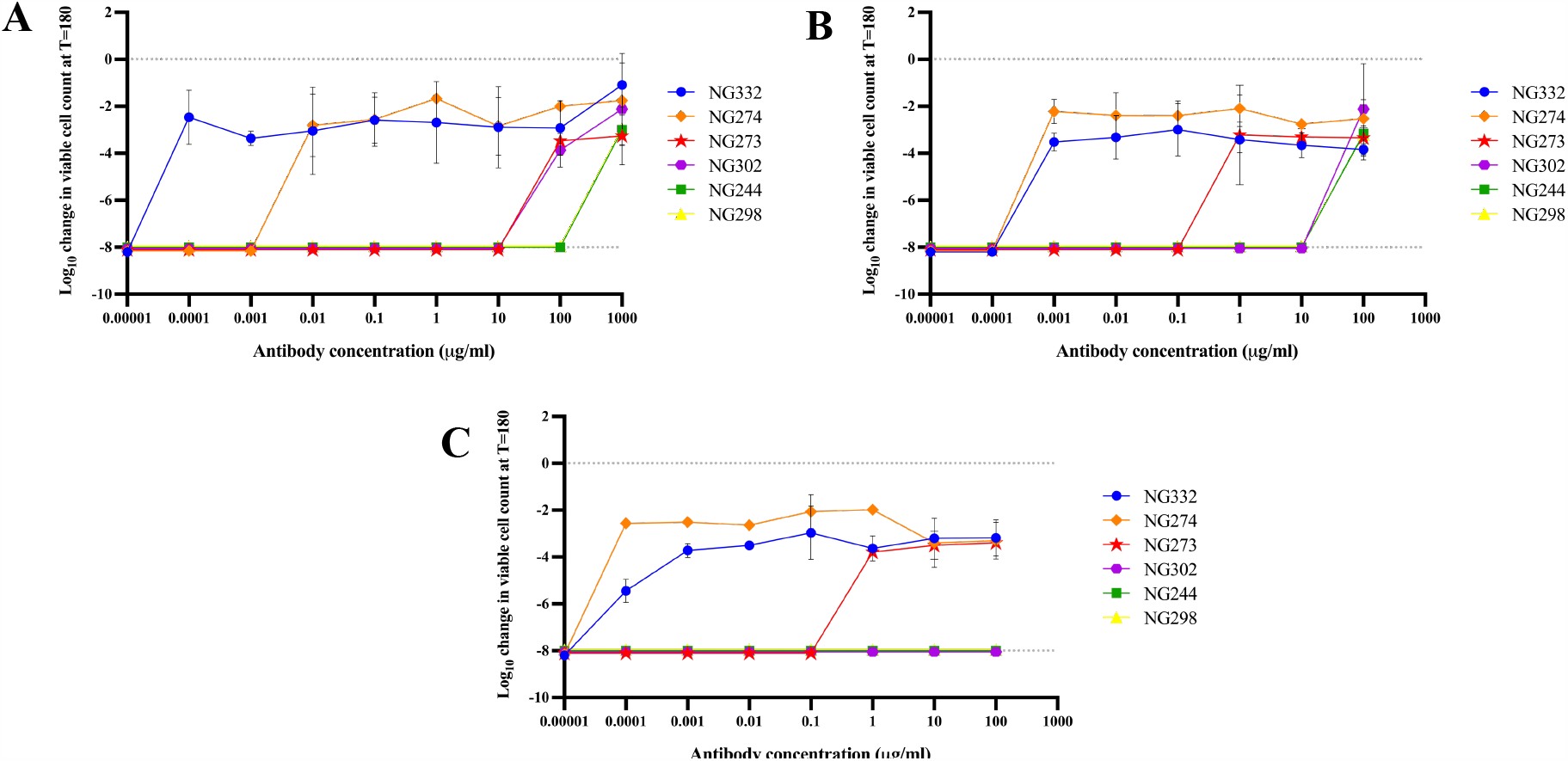
Effect of total IgG, IgM and IgA purified from inhibitory serum on autologous *Neisseria gonorrhoeae* isolates. A serum bactericidal assay was used to determine the effect of antibodies **A)** IgG, **B)** IgM and **C)** IgA purified from inhibitory serum on autologous clinical *N. gonorrhoeae* isolates. Cultures of isolates NG332, NG274, NG273, NG302, NG244 and NG298 were incubated with a mixture of 50% HCS and the indicated concentration of each autologous purified antibody. The change in viable colony forming units after 180 min was measured, with negative values corresponding to a decrease in viable bacteria when compared to the initial inoculum. Values at -8 indicate the limit of detection for samples from which no colonies formed. Data shown are the mean and standard deviation from two independent experiments.

To understand whether these inhibitory purified antibodies were able to initiate CMK, antibodies were also tested individually with 5% BRC as an exogenous complement source. Thus, these experiments tested the ability of the purified antibody isotypes to initiate CMK at varying concentrations. As expected, all isolates remained viable in 5% BRC alone (Supp. Fig. 2). These results showed that very low concentrations of IgG in all isolates were able to mediate CMK, as was IgM for all isolates, aside from NG273 and NG274, for which lower concentrations of antibody are required to be tested. IgA was not able to trigger CMK at any concentration, consistent with IgA being a poor inducer of complement activation. However, for most isolates tested, higher concentrations of IgG and IgM prevented CMK, mimicking the inhibitory phenotype of the autologous sera (Supp. Fig. 2).

Collectively, these data suggest that inhibition of HCS-mediated killing occurs at high concentrations with all isotypes tested, but IgG and IgM are able to initiate CMK at low concentrations in the presence of BRC. However, this inhibitory phenotype is most prominent using antibodies purified from NG274-serum and NG332-serum, suggesting that there might be isolate or antibody subclass-related differences driving this stronger phenotype.

### The effect of inhibitory serum on the survival of serum sensitive isolates

To investigate the mechanisms driving the ability of antibodies to prevent CMK, inhibitory NG332-serum was tested for its ability to kill serum-sensitive isolates NG233 and NG297. As expected, NG332-serum was not able to mediate CMK of its autologous isolate NG332, but it also failed to kill isolate NG233. However, inhibitory NG332-serum was able to mediate CMK of isolate NG297 (Fig. 4A). The same observation was made when using NG274-serum against these isolates (Fig. 4B). Conversely, both NG233-serum and NG297-serum were able to mediate CMK of both isolates NG332 and NG274 (Fig. 4C and D). To assess whether it was antibodies in the inhibitory NG332-serum and NG274-serum that protected NG233 from CMK, an SBA was set up as previously described using the IgG, IgM and IgA antibodies purified from the two inhibitory sera. Isolate NG233 was protected from CMK by the three antibody isotypes from both inhibitory NG332-serum and NG274-serum at all concentrations tested. However, these inhibitory antibodies were not able to protect isolate NG297 from CMK (data not shown). This raised the possibility that either differing surface antigens and/or different IgG subclasses are involved in the ability to block, or indeed, drive CMK.

**Figure 4.**
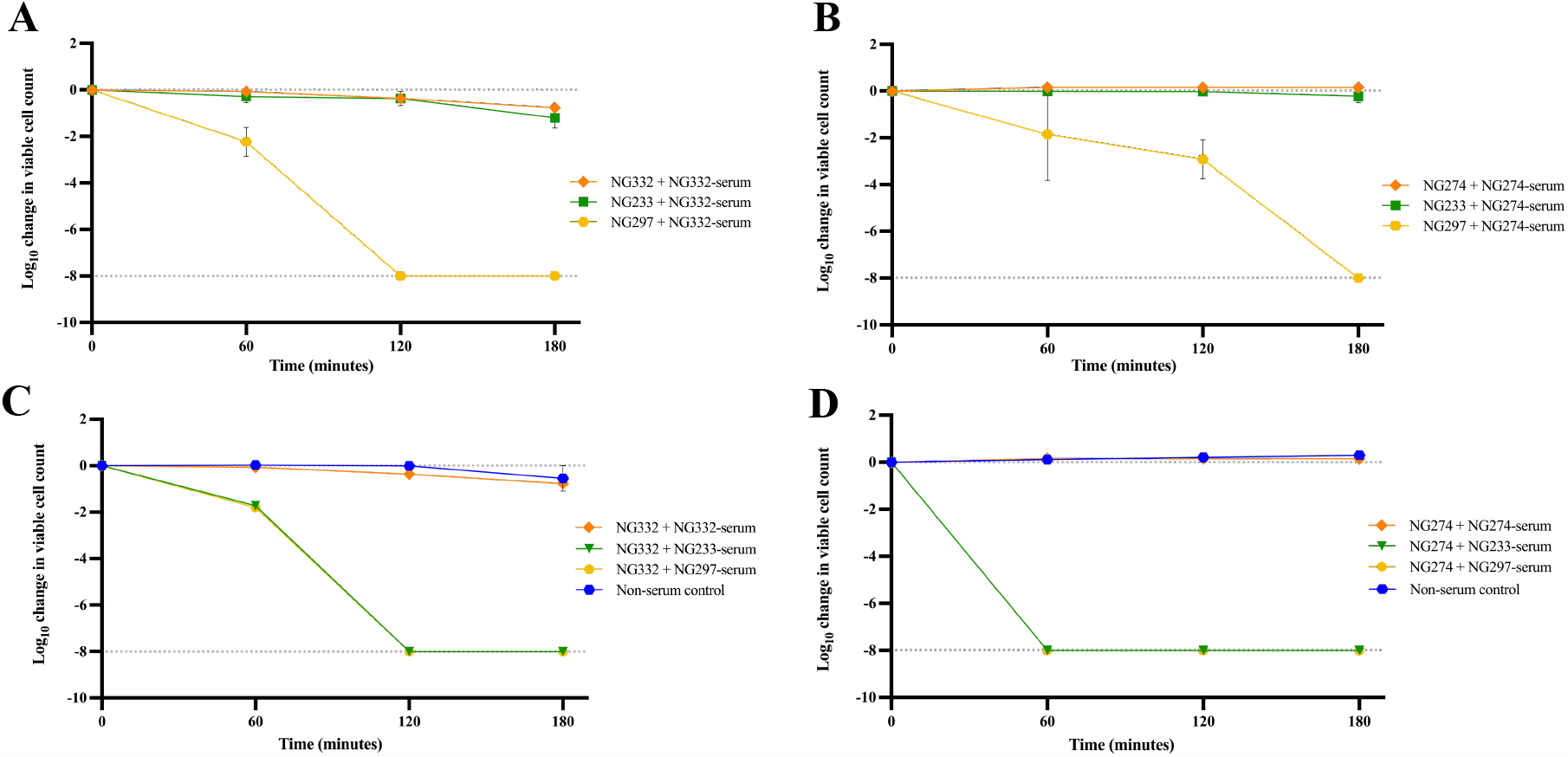
Serum bactericidal assays of inhibitory isolates NG332 and NG274 and their autologous inhibitory sera. A serum bactericidal assay was used to determine the sensitivity of isolates NG233 and NG297 in **A)** inhibitory NG332-serum and **B)** inhibitory NG274-serum. The susceptibility of inhibitory isolates **C)** NG332 and **D)** NG274 in both NG233-serum and NG297-serum were also assessed. The change in viable colony forming units after 60, 120 and 180 min were measured, with negative values corresponding to a decrease in viable bacteria when compared to the initial inoculum. Values at -8 indicate the limit of detection for samples from which no colonies formed. Data shown are the mean and standard deviation from two independent experiments.

### The effect of different IgG subclasses on the inhibition of complement-mediated killing by serum

Given the different effects of both NG332-serum and NG274-serum on the susceptibility of *N. gonorrhoeae* isolates to CMK, a whole cell ELISA was used to investigate binding titers of IgG subclasses to selected gonococcal isolates. First, the subclass of IgG binding to the eleven inhibitory candidates found to be resistant to the 1:1 serum mix were compared to an equal number of sensitive isolates with their matching sensitive serum, chosen to mirror the participant demographic of the resistant isolates. Inhibitory isolate NG319 was not assayed due to the lack of autologous serum, therefore only the remaining ten isolates were analysed. There were no significant differences in binding of total IgG, IgG1 and IgG4 between the resistant and sensitive isolates (Fig. 5A). This similarity between the total IgG binding to resistant and sensitive isolates was consistent with the flow cytometry data shown in Supp. Fig. 1. However, there was a slight increase in binding of IgG2 to resistant inhibitory isolates, but due to the limited number of isolates available, this was not statistically significant. There was a statistically significant decrease in binding of IgG3 in the inhibitory serum to resistant isolates compared to sensitive isolates (Fig. 5A). Moreover, the IgG2:IgG3 ratio bound to resistant isolates was increased compared with sensitive isolates (Fig. 5B), suggesting that the proportions of these antibodies binding to the bacterial cell surface may determine serum susceptibility.

**Figure 5.**
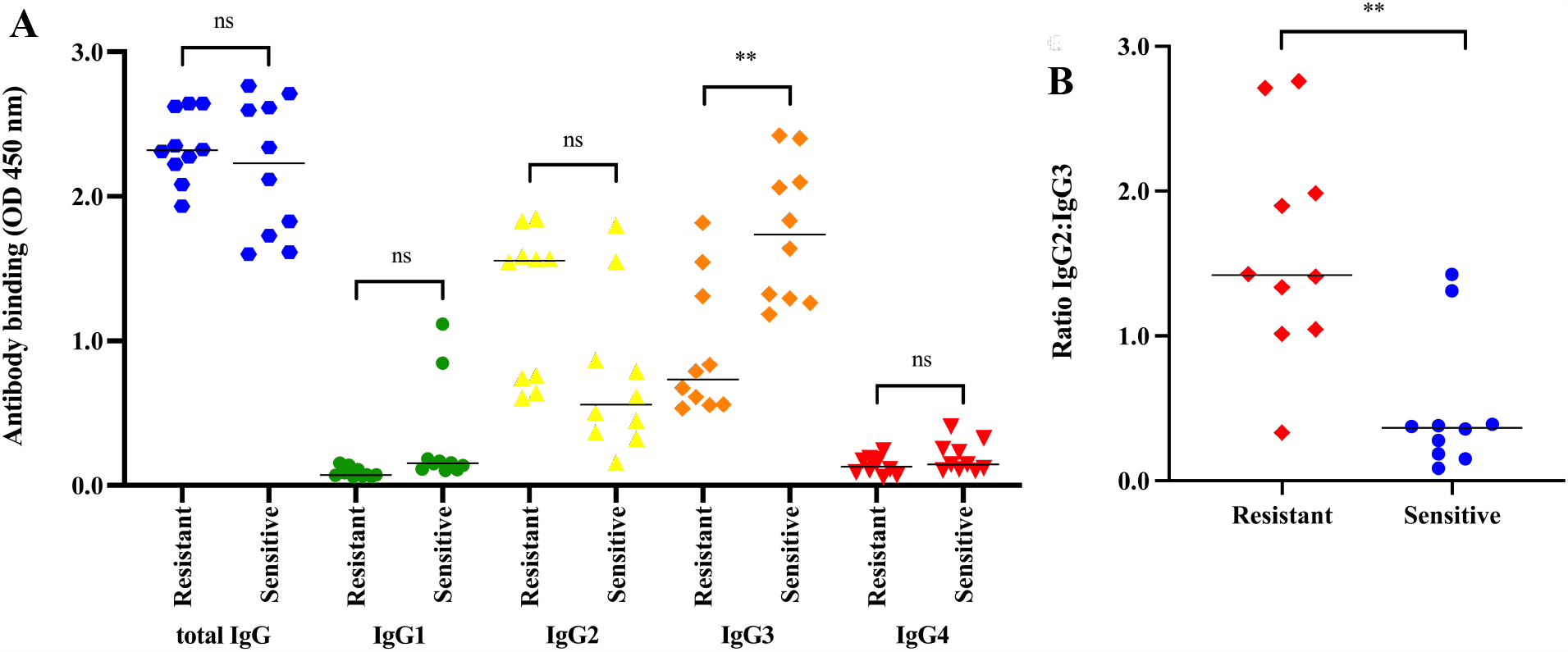
Comparison of IgG subclasses binding to serum-resistant and serum-sensitive isolates. An ELISA was used to investigate the amount of different IgG subclasses that bound to whole cell *N. gonorrhoeae*. **A)** Total IgG, IgG1, IgG2, IgG3 and IgG4 that bound to ten resistant isolates from autologous inhibitory serum were measured and compared to the binding of the same subclasses to ten isolates sensitive to autologous serum. **B)** The IgG2:IgG3 ratio that bound each of the ten resistant and sensitive isolates were calculated and compared. Data shown are the mean and standard deviation from two independent experiments. Statistical significance was measured using an unpaired t-test, where ** (*p*≤0.01).

We next wanted to determine whether differences in the IgG2:IgG3 ratio bound to the gonococcus could explain the differences in the susceptibilities of NG233 and NG297 to CMK by the inhibitory NG332-serum and NG274-serum shown in Fig. 4. Whole cell ELISAs revealed that there was no difference in binding of total IgG, IgG1 and IgG4 from either of the sera to these isolates (Fig. 6A-B). However, higher titres of IgG2 from NG332-serum were bound to isolates NG332, and NG233, which were protected from CMK, compared to NG297, which was sensitive to CMK (Fig. 6A). Conversely, lower titres of IgG3 were bound to NG332 and NG233, compared with NG297 (Fig. 6A). The same trend was observed when using NG274-serum against isolates NG274, NG233 and NG297 (Fig. 6B). These results reinforce that the IgG2:IgG3 ratio binding to the bacterial cell surface could be driving resistance to CMK. To test this theory further, binding of IgG subclasses from killing NG233-serum and NG297-serum to inhibitory isolates NG332 and NG274 were measured. There was no difference in binding of total IgG, IgG1 and IgG4 from any of the sera to isolate NG332 (Fig. 6C). However, there was a statistically significant decrease in binding of IgG2 to NG332 when incubated with killing NG233-serum and NG297-serum compared to its autologous inhibitory NG332-serum (Fig. 6C). There was also a statistically significant increase in binding of IgG3 from NG297-serum to isolate NG332 when compared to its autologous inhibitory serum, yet there was only a small increase in binding of IgG3 from NG233-serum. Similar results were obtained with inhibitory isolate NG274 (Fig. 6D). These data suggest that increased binding of IgG3 and decreased binding of IgG2 to the bacterium results in CMK of that isolate. They also suggest that the inhibitory antibody mechanism might depend on increased binding of IgG2 to the gonococcal cell surface. This could block bactericidal IgG3 from binding and initiating CMK.

**Figure 6.**
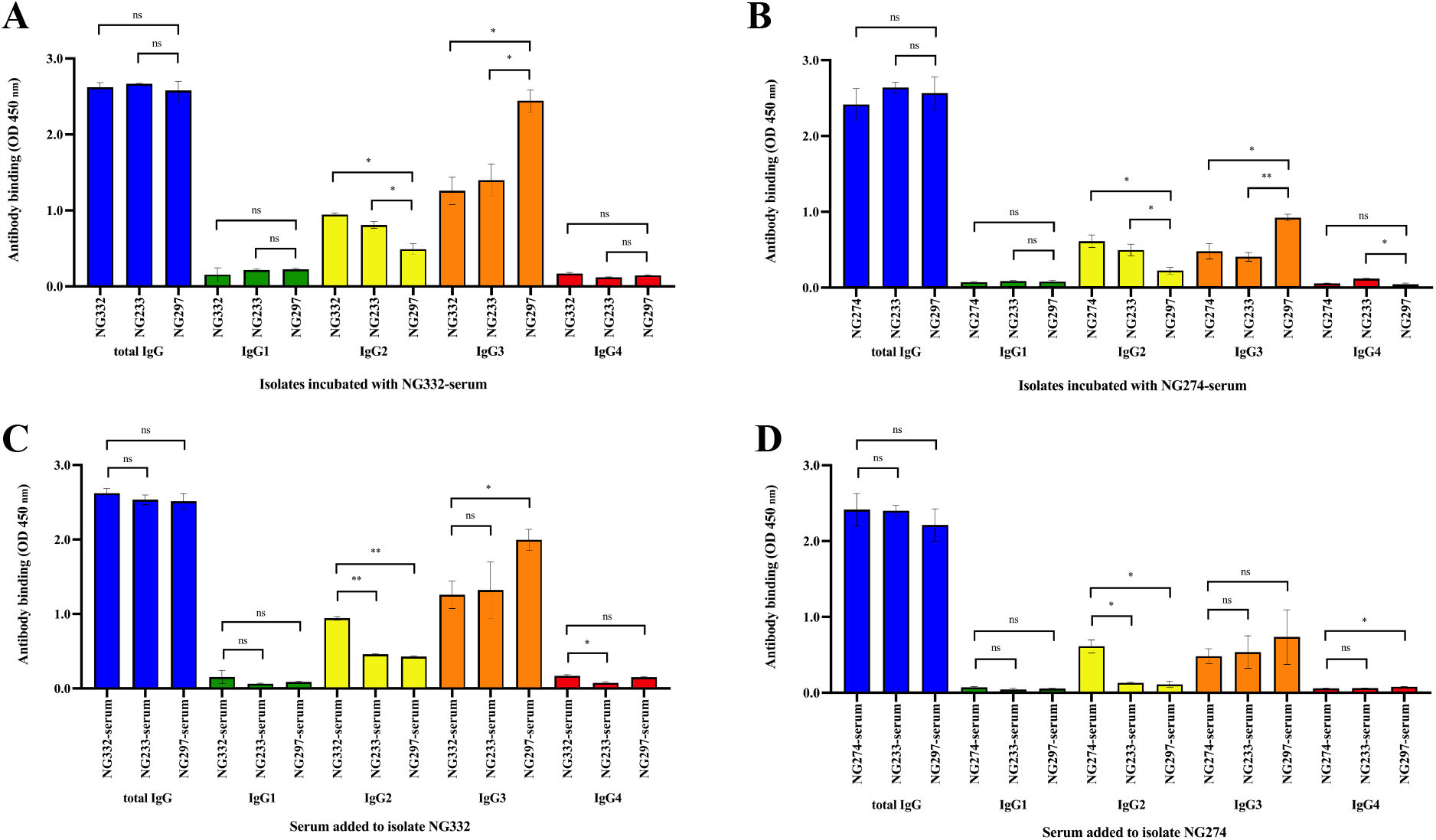
Comparison of binding of IgG subclasses from inhibitory sera to *Neisseria gonorrhoeae* isolates. An ELISA was used to investigate the amount of different IgG subclasses, including total IgG, IgG1, IgG2, IgG3 and IgG4, that bound to whole cell *N. gonorrhoeae* isolates from different sera. **A)** Binding to isolates NG332, NG233 and NG297 incubated with inhibitory NG332-serum and **B)** binding to isolates NG274, NG233 and NG297 incubated with inhibitory NG274-serum. **C)** Inhibitory NG332 incubated with autologous serum, NG233-serum and NG297-serum and **D)** inhibitory NG274 incubated with autologous serum, NG233-serum and NG297-serum. Data shown are the mean and standard deviation from two independent experiments. Statistical significance was measured using an unpaired t-test, where * (*p*≤0.05) and ** (*p*≤0.01).

## Discussion

Our investigation of serum susceptibility of clinical *N. gonorrhoeae* isolates from the G-ToG collection revealed that 3% (9/283) of participants recruited to the trial produced inhibitory antibodies. In contrast, Pham *et al*. (2021) and Wells *et al*. (2014) reported that one in three adults with cystic fibrosis and 20% of patients with non-cystic fibrosis bronchiectasis had inhibitory antibodies against *P. aeruginosa* infection, respectively. In addition, Coggon *et al*. (2018) found that 24% of patients with *E. coli* urosepsis had inhibitory antibodies. These data suggest that the prevalence of inhibitory antibodies during chronic infection is relatively high. In this study, however, the gonococcal infections were all acute and the incidence of inhibitory antibodies are relatively low. Whole genome sequencing and genome-wide association analyses of the collection of isolates did not show any significant genomic variants associated with resistance to autologous serum (data not shown). This indicates that the driver for this phenomenon could be at the transcriptional level in these isolates and/or mediated by fundamental differences in the host’s response to infection. Interestingly, we observed that participants with urethral infections and within the MSW sexual network were found to be significantly more likely to have protective sera. Thus, it would be of great interest to determine the clinical implications of inhibitory antibodies on recurrent gonorrhoea and patient outcomes in targeted follow up studies.

We report that IgG, IgM and IgA all have roles in the inhibition of CMK in a concentration-dependent manner. These data add to the body of evidence implicating antibodies IgG, IgM and IgA in driving the inhibition of CMK in response to Gram-negative bacterial infections in some individuals (MacLennan *et al*., 2010, Wells *et al*., 2014, Ray *et al*., 2011, Goh *et al*., 2016, Pham *et al*., 2021, Amir *et al*., 2005, Corbeil *et al*., 1988, Coggon *et al*., 2018). Broadly, the mechanism in which inhibitory antibodies block CMK in Gram-negative bacterial infections has not yet been characterised, although multiple theories have been proposed, as further detailed in a review by Torres *et al*. (2021). We show that an underlying mechanism of serum resistance for some gonococcal isolates could be related to an imbalance of the IgG2:IgG3 ratio binding to the bacterial cell surface. This is distinct from those previously described against other Gram-negative bacteria, where only very high titers of IgG2 bound to bacteria exhibit the inhibitory phenotype (MacLennan *et al*., 2010, Wells *et al*., 2014, Coggon *et al*., 2018). As mentioned, inhibitory antibodies against *N. gonorrhoeae* have previously been observed against the outer membrane protein RmpM against DGI strains (Rice *et al*., 1986). However, it is likely that the IgG2 identified in this study is targeting an alternative antigen, given the preference of IgG2 for binding to polysaccharide antigens (Vidarsson *et al*., 2014). Although outside the scope of this study, further work will characterise the antigens targeted by both the IgG2 and IgG3 antibodies identified in this study.

The cross-protection observed against *N. gonorrhoeae* by the OMV vaccine, MeNZB (Petousis-Harris *et al*., 2017), has motivated ambitions for development of a similar vaccine against gonorrhoea. Thus, the importance of the IgG2:IgG3 antibody ratios bound to the bacterial cell surface should be a consideration. Modification of *N. gonorrhoeae* vaccine strains such that their OMVs produce an increased amount of protein target for bactericidal IgG3 antibody and a decreased amount of target antigen for inhibitory IgG2 antibody might generate more efficient antibody-driven CMK, whilst limiting the production of inhibitory antibodies in vaccinated individuals. Many studies have suggested that a vaccine evoking a Th1 response in the host induces immune memory and infection clearance (Liu *et al*., 2013, Criss *et al*., 2009), unlike the Th17 response produced during natural infection (Gagliardi *et al*., 2011). Th1 development causes plasma cells to secrete high-affinity specific IgG3 antibodies (Li and Su, 2018). This is supported by the results observed in this study, where increased IgG3 binding correlated with bacterial killing, suggesting IgG3 binding might be important for protecting against gonococcal infection. Thus, antigens that stimulate this response might be crucial for vaccine-induced protection.

In summary, we have proposed a new mechanism to explain why a minority of participant antisera protect some gonococcal strains and, conversely why the majority kill this widespread pathogen. Clinical samples available in other laboratories should now be screened using the techniques described above to confirm or negate the proposed mechanism. The data obtained will not only reveal whether these inhibitory antibodies are predictive of clinical outcome and/or repeat infections but also provide a new focus for the development of an effective vaccine against gonorrhoea.

## Acknowledgments

We thank the patients that provided the samples used for this study from the G-ToG clinical trial. We would like to acknowledge the Clinical Immunology Service (CIS) staff, managed by Mr. Timothy Plant, who helped process the serum samples and for the use of the FACS Canto II. We thank Dr. Margaret Goodall (CIS) for kindly providing monoclonal IgG, IgM and IgA antibodies for flow cytometry. We acknowledge the Wellcome Trust Doctoral Training Programme in Antimicrobials and Antimicrobial Resistance to S.A.M. and A.E.R. (grant reference 108876/B/15/Z). We also acknowledge funding to J.D.C.R. from the Sexual Health Research Unit, University Hospital Birmingham NHS Foundation Trust.

## Author contributions

Conceptualisation: A.F.C., I.R.H., J.D.C.R. and A.E.R. Methodology: S.A.M., S.E.F., A.C. and N.K. Data analysis: S.A.M., J.A.C. and A.E.R. Writing: S.A.M., J.A.C. and A.E.R. Funding acquisition: I.R.H., J.D.C.R. and A.E.R.

---

**Supplementary Figure 1.**
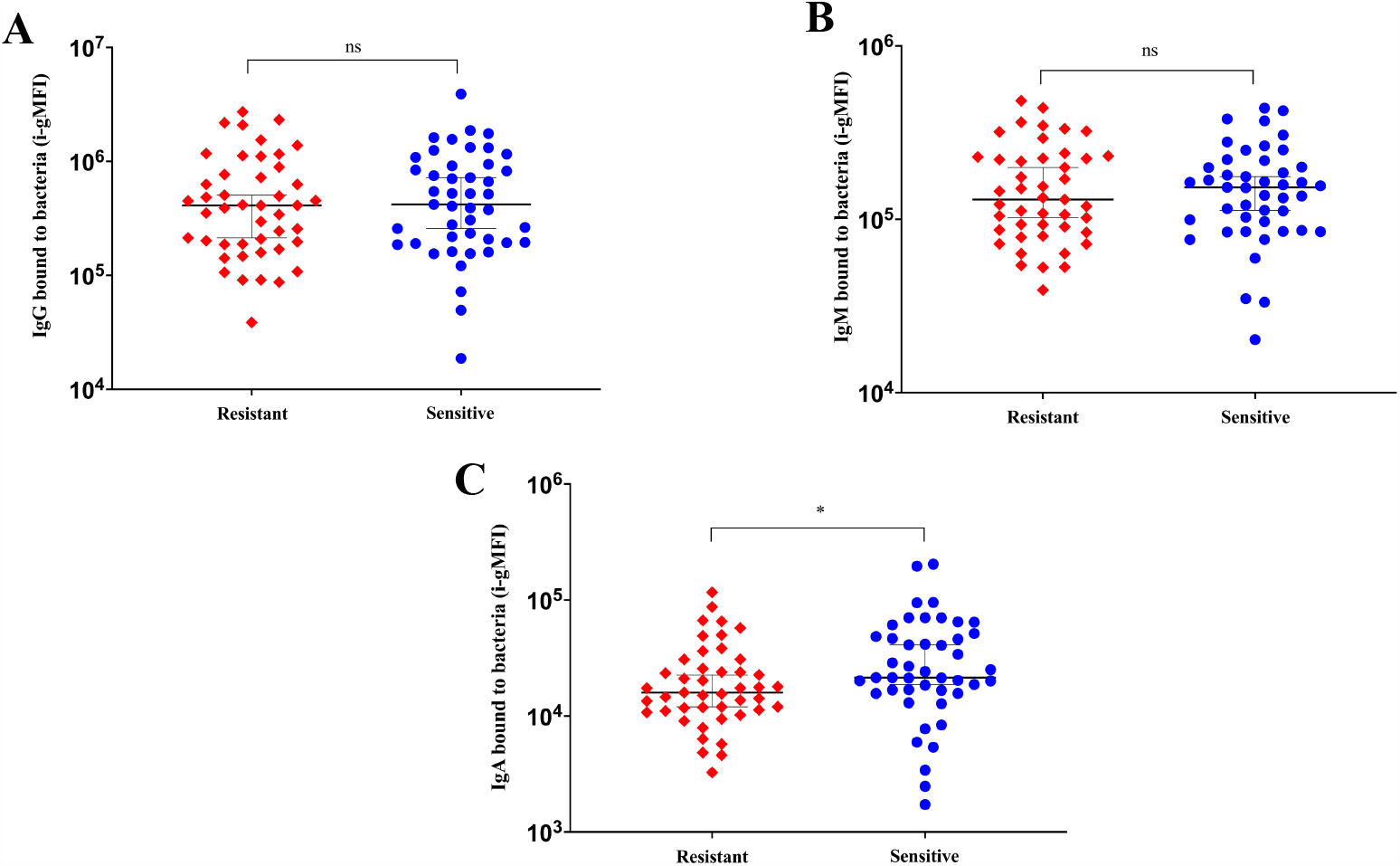
Antibody binding to resistant and sensitive isolates. A flow cytometric assay was used to analyse antibody binding to the 46 isolates resistant to autologous serum, and 46 isolates sensitive to autologous serum. Cultures of each isolate were incubated with autologous serum and binding of **A)** IgG, **B)** IgM and **C)** IgA were measured by flow cytometry using fluorescently labelled secondary antibodies. Shown is the integrated geometric mean fluorescence intensity (i-gMFI) for each serum protein and all of the 92 isolates, with isolates resistant to the 1:1 serum mixture highlighted in yellow. The median for each group was calculated and a 95% confidence interval (CI) was applied. Statistical significance was measured using a Mann-Whitney test, where *p*≤0.05 (*).

**Supplementary Figure 2.**
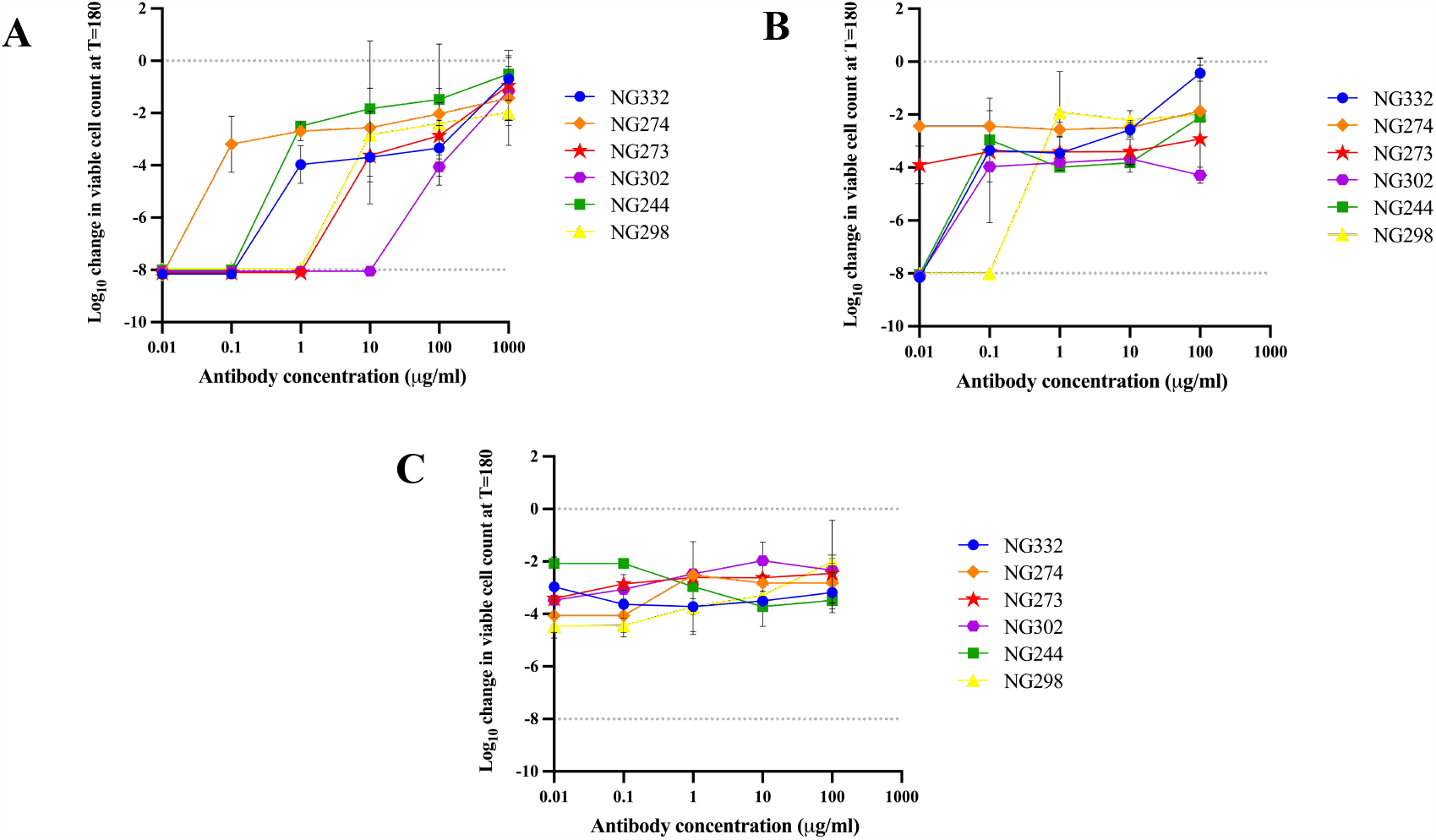
Effect of total IgG, IgM and IgA purified from inhibitory serum on autologous *Neisseria gonorrhoeae* isolates in the presence of baby rabbit complement alone. A serum bactericidal assay was used to determine the effect of antibodies **A)** IgG, **B)** IgM and **C)** IgA purified from inhibitory serum on the matching *N. gonorrhoeae* isolates. Cultures of isolates NG332, NG274, NG273, NG302, NG244 and NG298 were incubated with a mixture of 5% BRC and each autologous purified antibody at the indicated concentrations. The change in viable colony forming units after 180 min was measured, with negative values corresponding to a decrease in viable bacteria when compared to the initial inoculum. Values at -8 indicate the limit of detection for samples from which no colonies formed. Data shown are the mean and standard deviation from two independent experiments.

**Supplementary Table 1.** Comparisons of clinical metadata between the whole population and participants with inhibitory sera. Proportions of participants recruited to the G-ToG clinical trial in the indicated categories amongst the whole population (n=283) versus the participants with inhibitory sera (n=9). Statistical significance was calculated using a 2-tailed z-test. Significant values (*p*<0.05) are shown in red bold text.

| Category | Subcategory | Total population (n=283) | Total population (%) | Inhibitory population (n=9) | Inhibitory population (%) | 2-proportional z-test score | P value |
| --- | --- | --- | --- | --- | --- | --- | --- |
| Geographical location | London | 90 | 31.8 | 4 | 44.4 | -0.799 | 0.212 |
|  | Outside London | 192 | 67.8 | 5 | 55.6 | 0.775 | 0.219 |
|  | Unknown | 1 | 0.4 | 0 | 0.0 | 0.179 | 0.429 |
| Ethnicity | Black | 36 | 12.7 | 1 | 11.1 | 0.143 | 0.443 |
|  | Asian | 19 | 6.7 | 1 | 11.1 | -0.514 | 0.304 |
|  | Other ethnic group | 13 | 4.6 | 1 | 11.1 | -0.901 | 0.184 |
|  | Mixed ethnic group | 23 | 8.1 | 0 | 0.0 | 0.891 | 0.186 |
|  | White | 192 | 67.8 | 6 | 66.7 | 0.074 | 0.470 |
| Gender | Female | 51 | 18.0 | 1 | 11.1 | 0.533 | 0.297 |
|  | Male | 231 | 81.6 | 8 | 88.9 | -0.557 | 0.289 |
|  | Other | 1 | 0.4 | 0 | 0.0 | 0.179 | 0.429 |
| Sexual network | WSM | 51 | 18.0 | 1 | 11.1 | 0.533 | 0.297 |
|  | MSW | 59 | 20.8 | 4 | 44.4 | -1.694 | **0.045** |
|  | MSM | 172 | 60.8 | 4 | 44.4 | 0.986 | 0.162 |
|  | Other | 1 | 0.4 | 0 | 0.0 | 0.179 | 0.429 |
| Site of infection | Cervical | 40 | 14.1 | 1 | 11.1 | 0.257 | 0.399 |
|  | Rectal | 80 | 28.3 | 2 | 22.2 | 0.397 | 0.346 |
|  | Pharyngeal | 49 | 17.3 | 0 | 0.0 | 1.368 | 0.086 |
|  | Urethral | 156 | 55.1 | 8 | 88.9 | -2.010 | **0.022** |
|  | Unknown | 11 | 3.9 | 0 | 0.0 | 0.603 | 0.273 |
| Previous gonorrhoea infections | Previous infection | 117 | 41.3 | 5 | 55.6 | -0.851 | 0.197 |
|  | No previous infections | 164 | 58.0 | 4 | 44.4 | 0.807 | 0.210 |
|  | Unknown | 2 | 0.7 | 0 | 0.0 | 0.253 | 0.400 |
| HIV status | Positive | 43 | 15.2 | 2 | 22.2 | -0.575 | 0.283 |
|  | Negative | 235 | 83.0 | 7 | 77.8 | 0.412 | 0.340 |
|  | Unknown | 5 | 1.8 | 0 | 0.0 | 0.402 | 0.344 |
| Symptoms | No symptoms | 109 | 38.5 | 2 | 22.2 | 0.991 | 0.161 |
|  | Symptoms | 174 | 61.5 | 7 | 77.8 | -0.991 | 0.161 |

